# Anatomy of the heart of the leatherback turtle

**DOI:** 10.1101/2022.02.22.481434

**Authors:** Bjarke Jensen, Henrik Lauridsen, Grahame Webb, Tobias Wang

**Author notes:** corresponding author: Bjarke Jensen, Ph.D. Amsterdam UMC, Department of Medical Biology, Room L2-106, Meibergdreef 15, 1105AZ Amsterdam, The Netherlands, Fax: not available.

## Abstract

Non-crocodylian reptiles have hearts with a single ventricle, which is partially separated by a muscular ridge that provide some separation of blood flows. An exceptional situation exists in monitor lizards and pythons, where the ventricular left side generates a much higher systolic blood pressure than the right side, thus resembling mammals and birds. This functional division of the ventricle depends on a large muscular ridge and may relate to high metabolic demand. The large leatherback turtle (<1000 kg), with its active ocean-going lifestyle and elevated body temperatures, may have similar adaptations. Here, we report on the anatomy the hearts of two leatherback turtles. One stranded in Ballum, Denmark in 2020, and was examined in detail, supplemented by observations and photos of an additional stranding specimen from Canada. The external morphology of the leatherback heart resembles that of other turtles, but it is large. We made morphometric measurements of the Ballum heart and created an interactive 3D model using high resolution MRI. The volume of the ventricle was 950 ml, from a turtle of 300 kg, which is almost twice as large as in other reptiles. The Ballum heart was compared to MRI scans of the hearts of a tortoise, a python, and a monitor lizard. Internally, the leatherback heart is typical of non-crocodylian reptiles, and did not contain the well-developed septation found in pythons and monitor lizards. We conclude that if leatherback turtles have exceptional circulation needs, they are sustained with a relatively large but otherwise typical non-crocodylian reptile heart.

## Introduction

The septation of the cardiac ventricle in non-avian reptiles is much more variable than in other vertebrate groups (Poelmann & Gittenberger-de Groot, 2019; Jensen & Christoffels, 2020), and this variation may well shed light on the evolution of the two ventricles in mammals and birds, which both evolved from an ancestor with a single ventricle (Kardong, 2006; Jensen et al., 2010a; Poelmann *et al*., 2014; Katano *et al*., 2019).

Brücke (1852) first established the heart of non-crocodylian reptiles consists of two atria and a single ventricle (Brücke, 1852; Webb, 1979). Later studies clarified that reptiles have an additional chamber because the venous sac behind the right atrium, the sinus venosus, contracts well before the atria and aids the filling of the right atrium (Johansen, 1959; Valentinuzzi & Hoff 1972; Jensen *et al*., 2017). Brücke also established that the single ventricle is partially septated into three major compartments: i) The right-most is the cavum pulmonale that leads to pulmonary artery (see label ‘CP’ in Figure 1), ii) The left-most compartment (cavum arteriosum, CA), which receives the inflow from the left atrium, and iii) the smaller cavum venosum, situated between the CP and the CA, that receives the inflow from the right atrium. Two bi-leaflet valves guard the outflows into the left and right aortas in the cranial part of cavum venosum, and oxygen-rich blood from the left-situated cavum arteriosum therefore must pass the through the cavum venosum during cardiac systole before being ejected into the systemic arteries. The right-situated cavum pulmonale receives blood from the right atrium in diastole, which also passes through the cavum venosum. The cavum venosum is thus a cross-road of filling in diastole and ejection in systole, and hence an important site of intracardiac shunting (Hicks, 1998; Burggren *et al*., 2020).

**Figure 1.**
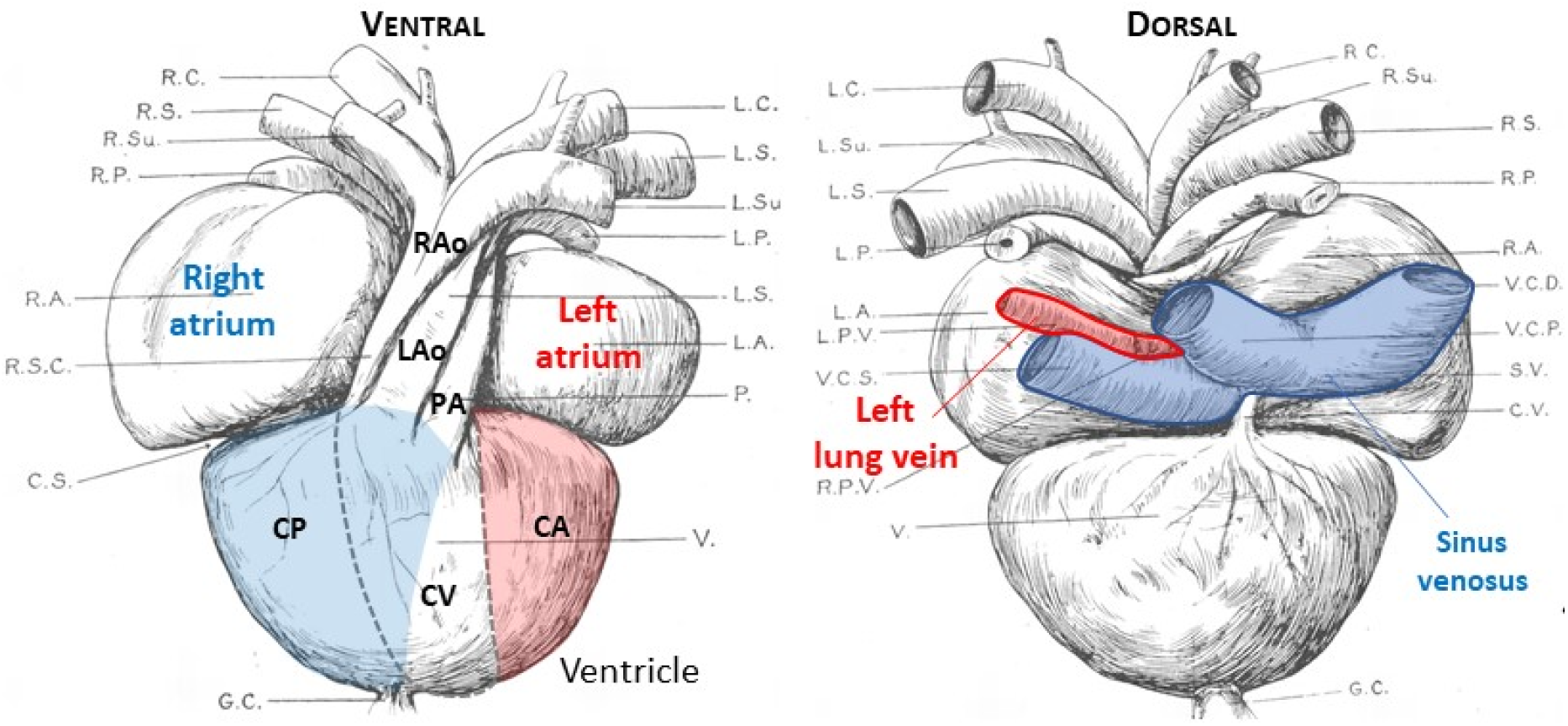
External view of the heart of a leatherback turtle. In the ventricle, the red color indicates the cavum arteriosum (CA) and blue indicates the cavum pulmonale (CP). Notice CP is partly ventral to the cavum venosum (CV), which is indicated by the blue area within the dashed-line perimeter of the CV. LAo, left aorta; L.P.V. (shaded red) is left lung vein; PA; pulmonary artery; RAo, right aorta. Adapted from O’Donoghue (1918). The original abbreviations are; C.S., coronary sulcus; C.V., coronary vein; G.C., gubernaculum cordis; L.A., left auricle; L.C., left carotid artery; L.P., left pulmonary artery; L.P.V., left pulmonary vein; L.S., left systemic arch; L.Su., left subclavian artery; P., pulmonary arch; P.V., pulmonary vein; R.A., right auricle; RA., right carotid artery; R.P., right pulmonary artery; R.P.V., right pulmonary vein; R.S., right systemic artery; R.S.C., right systemico-carotid trunk; R.Su., right subclavian artery; S.V., sinus venosus; V., ventricle; V. C.D., right pre-caval vein; V.C.P., post. caval vein; V.C.S., left pre-caval vein.

The cavum venosum is small in monitor lizards and particularly in pythons (Webb *et al*., 1971; Jensen *et al*., 2014). This reflects, at least in part, that the ventricular muscle underneath the atrioventricular valve is particularly prominent. This structure, which separates the cavum arteriosum from the cavum venosum, is called the vertical septum. A second septum, called the muscular ridge, is also more developed in monitors and pythons than in other reptiles, and further confine the volume of the cavum venosum. Finally, across from the muscular ridge is the bulbuslamelle (Greil, 1903), which is so developed in monitors and pythons that it together with the muscular ridge, can form a pressure-tight seal in systole. This shields the low pressure of the cavum pulmonale and pulmonary circulation from the almost mammal-like higher pressure generated in the cavum arteriosum and cavum venosum (White, 1968; Burggren & Johansen, 1982; Wang *et al*., 2003; Jensen *et al*., 2010a). The presence of this substantial septation is revealed on the ventricular surface by the presence of large coronary vessels, similar to the setting in crocodylians, birds, and mammals (Jensen *et al*., 2014).

Not much is known of the cardiac anatomy and physiology of the leatherback sea turtle (*Dermochelys coriacea*). The most detailed anatomical study is more than a century old (O’Donoghue, 1918) and reports that the leatherback heart is configured much like other reptile hearts (O’Donoghue, 1918). The external appearance is also much akin to other chelonians (Wyneken, 2001) (Figure 1). Later, Adams (1962) reported large foramens between the major arteries in a single stranded leatherback. Adams (1962) also described numerous dimples, or under-developed foramens, on the luminal side of these vessels, and speculated that the foramens are a common feature of the leatherback heart. The leatherback ventricle has also been noted as having “an exceptionally well-developed muscular ridge” (Wyneken, 2009), although no visual or other supporting evidence was provided. Thus, the degree to which the muscular ridge compares with well-developed one of monitors and pythons is unclear. When these observations are considered in relation to the large body size, the capacity for long oceanic migrations, and the ability to maintain a body temperature several degrees higher than the surrounding sea water (Paladino *et al*., 1990; James & Mrosovsky, 2004), leatherbacks may be another reptile in which the ventricle has adapted to better exploit pressure separation.

Here we undertook an anatomical study of the hearts of two large leatherback turtles to assess whether the leatherback heart is typical of chelonians or has features similar to monitors and pythons. Should the latter be the case, it is likely that the leatherback ventricle has pressure separation which is considered a prerequisite for high metabolic rates (Quick & Ruben, 2009; Lovegrove, 2019; Grigg *et al*., 2021).

## Materials and Methods

### Animals

The reported findings are mostly based on the heart of the leatherback that stranded near Ballum (Denmark) on 3^rd^ of November 2020. It had a curved carapace length of 148 cm and was estimated to weigh 300 kg. It was brought to The Fisheries and Maritime Museum in Esbjerg, Denmark, given the specimen ID “R2117”, and kept frozen until its dissection on 6^th^ of August 2021. The other heart examined came from a stranded leatherback in Canada (2009), which was examined in Australia, mainly to confirm the foramen between the arterial and pulmonary arteries described by Adams (1962). Photos taken during the dissection (Sept 2011) allowed verification of observations made on the Ballum heart. It is stated explicitly in the Results when findings on the Ballum heart could be verified in the Canadian heart. Comparisons are made between images from MRI of hearts of a *Geochelone* tortoise and a Burmese python (adapted from Jensen *et al*., 2014) as well as an Asian water monitor lizard (adapted from Hanemaaijer *et al*., 2019).

### Dissection and fixation

The Ballum heart was dissected free and all major structures were more or less intact. Accordingly, the dissected tissue included parts of the major arteries, esophagus, and the left lung, together with coagulated blood, and weighed in total 11.2 kg. This specimen was kept in a 25 l bucket filled with 4% formaldehyde buffered to pH 7.4. The formaldehyde was renewed after three days and total immersion fixation time was one week. After this, the heart was transferred to a phosphate buffered saline and washed twice for three days to remove residual formaldehyde in preparation for magnetic resonance and x-ray computed tomography imaging (MRI and CT).

### Magnetic resonance and x-ray computed tomography imaging and 3D reconstruction

Magnetic resonance imaging (MRI) was performed on an Siemens Magnetom Skyra system equipped with a Flex Large 4 surface coil using a T2-weighted 3D sequence with the following parameters: field strength = 3 T, repetition time = 1000 ms, echo time = 132 ms, flip angle = 120°, field-of-view = 288 × 288 × 160 mm^3^, spatial resolution 0.5 mm isotropic, number of averages = 4, acquisition time = 3.7 h.

X-ray computed tomography (CT) was performed using a Toshiba Aquillon Prime SP system with the following parameters: x-ray tube voltage = 120 kVp, x-ray tube current = 250 mA, integration time = 1000 ms, field-of-view = 410.1 × 410.1 × 404.0 mm^3^, spatial resolution = 0.8 mm isotropic, convolution kernel = FC18, acquisition time = 90 s.

The MRI and CT generated image stacks were imported to the 3D software Amira (version 3D 2021.2, FEI SAS, Thermo Fisher Scientific). Labeling of structures were done in the Segmentation Editor module. The volumes of labelled structures were retrieved using the Materials Statistics module. Distances were measured in Orthographic view using the Measure Line tool on either images of the image stack or on a volume rendering of the label file. The Amira model was converted to an interactive 3D pdf as previously described (de Bakker *et al*., 2016).

## Results

Figure 2A shows the ventral side of the heart *in situ* of the Ballum leatherback. The coloration of the ventricle suggested that the tissue had not decomposed to any great extent. Numerous coronary vessels were found on the ventricular surface, but not in a similar configuration as in pythons, monitors and crocodylians, where the major coronary vessels run along the surface of major septation. Overall, the proportions and shapes of the cardiac chambers corresponded well to the previous accounts of leatherback hearts. A pronounced guberculum cordis extended from the ventricular apex. Figure 2B exemplifies one of the MR generated images. Larger structures such as the atrial septum can be identified. Numerous straight lines were observed in the cavities of the chambers and vessels on both MRI and CT images. We suspect these lines reflect structured coagulation of the red blood cells and they were not perceived as tissue. Also, thick somewhat concentric bands are seen in the ventricular muscle. These bands were not obvious on CT (Fig. 2B), which suggest that they do not reflect substantial differences in tissue density and we believe they are artefacts of fixation. From the MRI image stack, we reconstructed the chambers, valves at the chamber junctions, and the major arteries (Fig. 2C). The interactive pdf of this model, in which each labelled structure can be made visible or transparent, can be found in the supplement (Supplementary Figure 1). The ventricular wall (including septums), which was the best demarcated wall of the four chambers, comprised 950 ml. Concerning the atria, the atrial septum was 29 ml, the right atrial wall was 289 ml, and the left atrial wall was 125 ml. The wall of the sinus venosus was extremely faint in many images and it could not be reconstructed such that the tissue volume could be estimated. The estimated ventricular index of 0.33 % (950 ml / 300 kg) is quite high for a reptile where the average ventricular index is approximately 0.2% (Jensen et al 2014). The superficial appearance of the leatherback heart is typically chelonian and it resembles less that of the squamate monitors and pythons (Fig. 2C).

**Figure 2.**
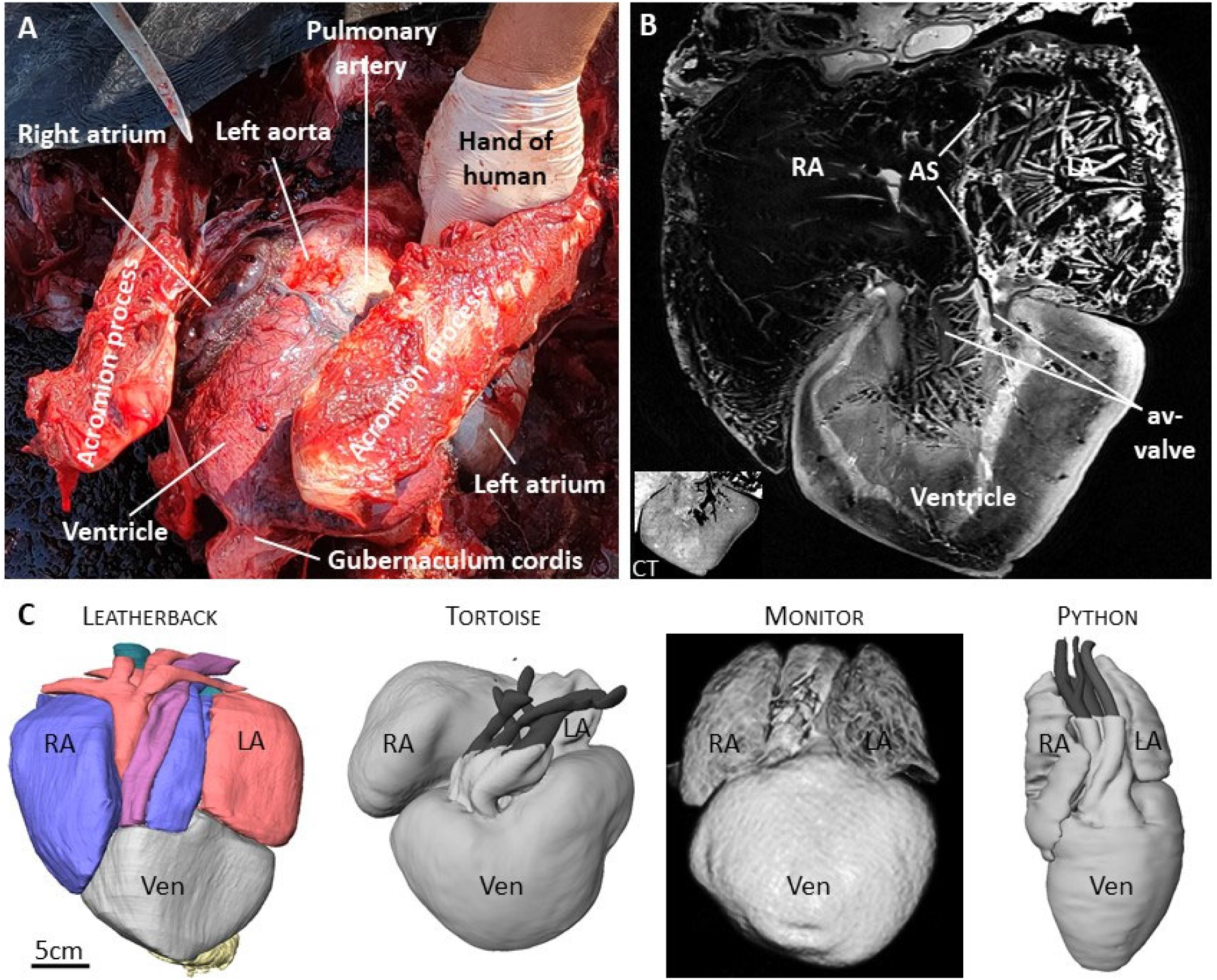
Heart of the Ballum turtle. **A**. Caudal-ventral in-situ view of the heart of the Ballum turtle. **B**. Example of MR-generated image of the heart. In the ventricle several bands are seen, but these are likely artefacts because the bands are not detected on CT (insert on the bottom left). **C**. Reconstruction of the Ballum turtle heart, next to reconstructions based on MRI of the hearts of the red-footed tortoise, the Asian water monitor, and the Burmese python.

The three sinus horns became confluent dorsal to the sinuatrial junction (Fig. 2C). While the sinuatrial junction was not wide in proportion to the right atrium, it was still well beyond 2 cm in width (Fig. 3A). The junction was guarded by two leaflets as is typical in reptiles (Fig. 3A). A large right leaflet and a smaller left leaflet guarded the right and left atrioventricular opening respectively. These leaflets were merged medially to each other and to the atrial septum, as is typical in non-crocodylian reptiles (Fig. 3B). The right and left atrioventricular openings were dorsal and to the left of the aortas. In the Canadian specimen, when the apical region of the ventricle had been resected, a similar atrioventricular valve was found (Fig. 3C D). No pronounced septation was found under the atrioventricular valve of the Canadian heart (Fig. 3D).

**Figure 3.**
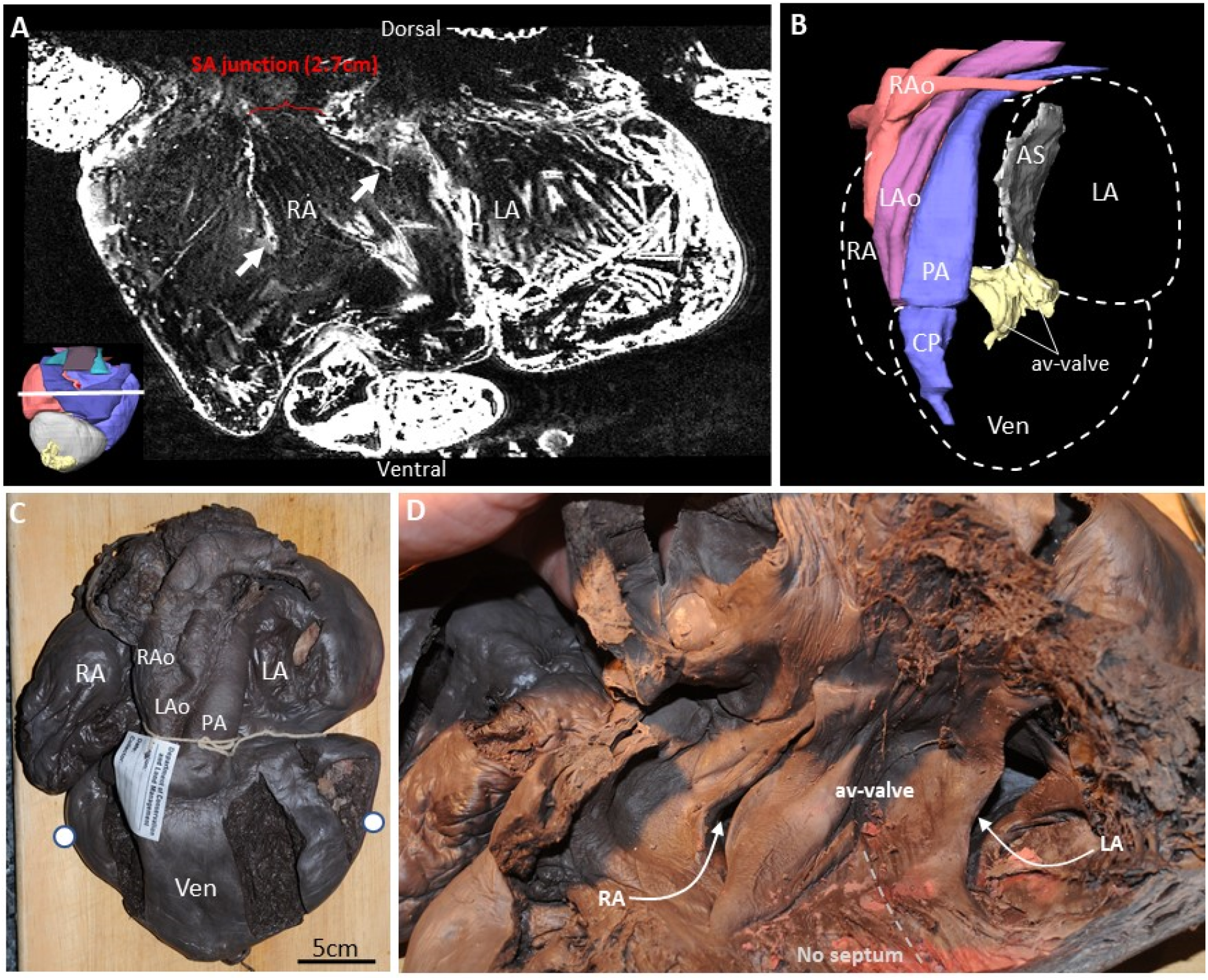
Chamber junction valves. **A**. The atria in the transverse plane, showing the two leaflets (white arrows) of the sinuatrial valve at the base of which is the sinuatrial junction which at this location 2.7cm wide. In the insert on the bottom-left, the section plane is indicated by a white line. **B**. Reconstruction of the atrioventricular valve (av-valve) showing that it is connected to the atrial septum (AS), and it is at some distance from the cavum pulmonale (CP) which leads to the pulmonary artery (PA). **C**. Ventral view of the Canadian heart. The two white dots on the ventricle (Ven) indicates the plane of cutting used to produce the view in D. **D**. When the apical part is removed and the ventral part of the ventricle is lifted, the av-valve is exposed showing two large leaflets, one for each of the atrial inflows. A substantial communication is possible between the left and right side of the ventricle because there is no ventricular septum immediately below the av-valve, leading some authors to consider this part as a single chamber, the cavum dorsale. LA, left atrium; LAo, left aorta; RA, right atrium; RAo, right aorta.

Images from MR revealed the absence of pronounced septation under the atrioventricular valve in the Ballum heart (Fig. 4A). Dorsally and ventrally, the atrioventricular valve was embedded in ventricular myocardium, as is typical in non-crocodylian reptiles (Fig. 4B). In this way, both the Ballum and Canadian hearts resembled the heart of other chelonians (Fig. 4C). The ventricular muscle in monitors and pythons, on the other hand, reaches much closer to the atrioventricular valve and is referred to as the vertical septum (Fig. 4D-E).

**Figure 4.**
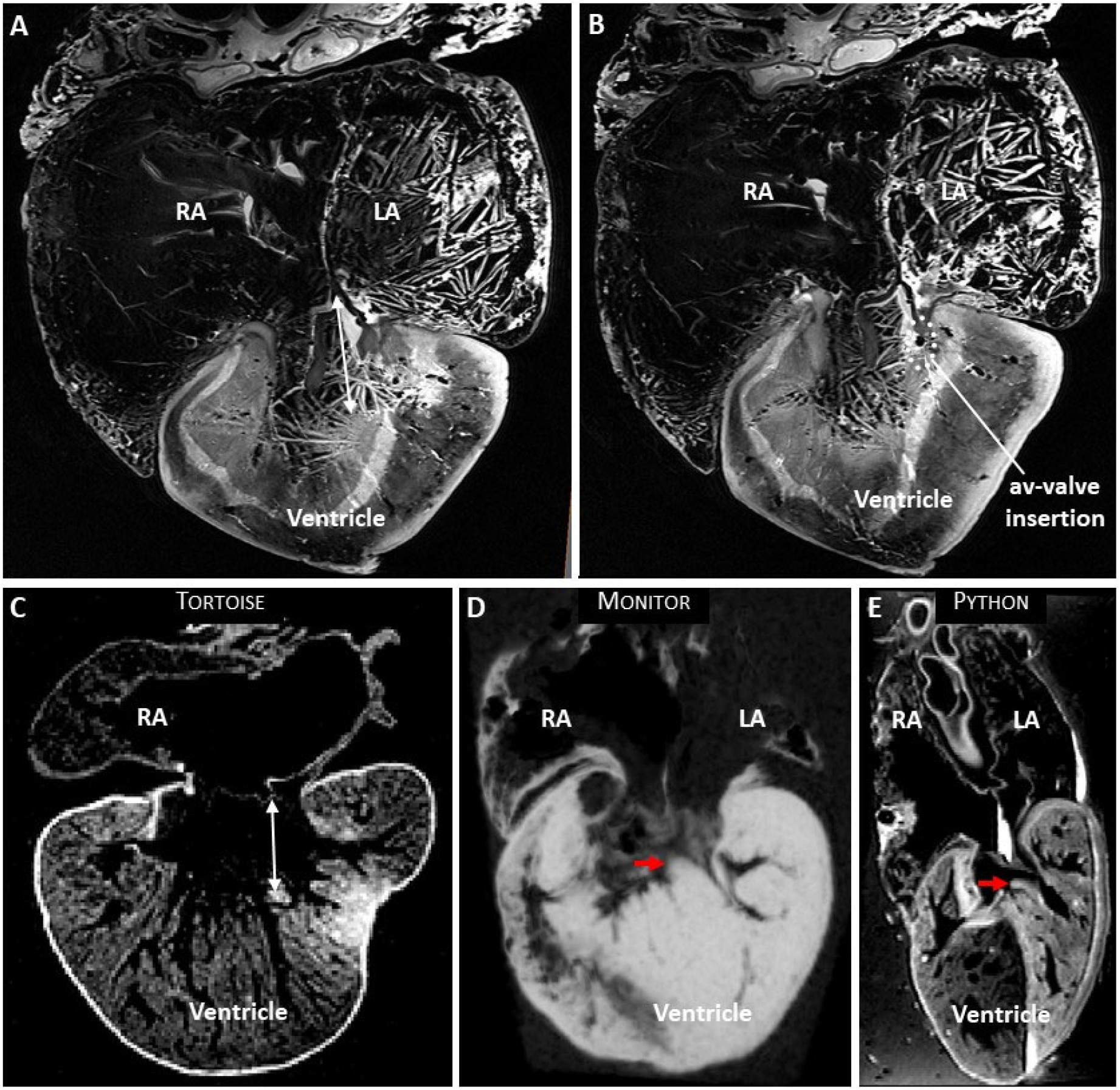
The vertical septum of the ventricle. **A**. In the leatherback ventricle, a large gap (double-headed arrow) is found between the ventricular muscle and the point where the two leaflets of the av-valve connects to the atrial septum. The shown image is approximately in the horizontal plane. **B**. Ventrally (and dorsally) the av-valve is inserted in ventricular muscle, an example of this is indicated with the dashed ovale. **C**. The tortoise also has a large gap between the ventricular muscle and the av-valve. **D**-**E**. In the monitor and python, the vertical septum (red arrow) almost makes contact with the av-valve. LA, left atrium; RA, right atrium.

The most pronounced septation of the Ballum ventricle was the muscular ridge (Fig. 5A). It harbors the base of the ventral leaflet of the valve of the left and right aorta and the dorsal leaflet of the pulmonary artery. Opposite to it is the bulbuslamelle, which harbors the dorsal leaflet of the valve of the left and right aorta. As is typical of non-crocodylian hearts, the muscular ridge is close to the horizontal plane near the arterial valves (Fig. 5A). It then has a helical course towards the apex, such that its free margin makes contact to do the dorsal-right wall of the ventricle (Fig. 5B). At least in the Canadian heart, there was a substantial space between the free edge of the muscular ridge and the opposite wall (Fig. 5C). This configuration is similar to that of the tortoise (Fig. 5D), and does not resemble the python or monitor hearts where the muscular ridge is much larger (Fig. 5D-E). It is not unusual for reptiles and turtles in particular to have a substantial cartilaginous deposit where the muscular ridge merges with the truncus (Poelmann *et al*., 2017) but we could not detect such deposit on the CT or MRI (data not shown). The bulbuslamelle is also very developed in monitors and pythons, but this is best appreciated in ventricles that are not fully contracted, such as in the python heart depicted in Figure 5F. Notice that in the python heart, there is only a narrow gap between the muscular ridge and the bulbuslamelle even in the dilated state, which is in contrast to that of chelonians (Fig. 5C-D). Due to the state of contraction in the Ballum heart, however, we could not assess whether the bulbuslamelle was typical or more like that of monitors and pythons.

**Figure 5.**
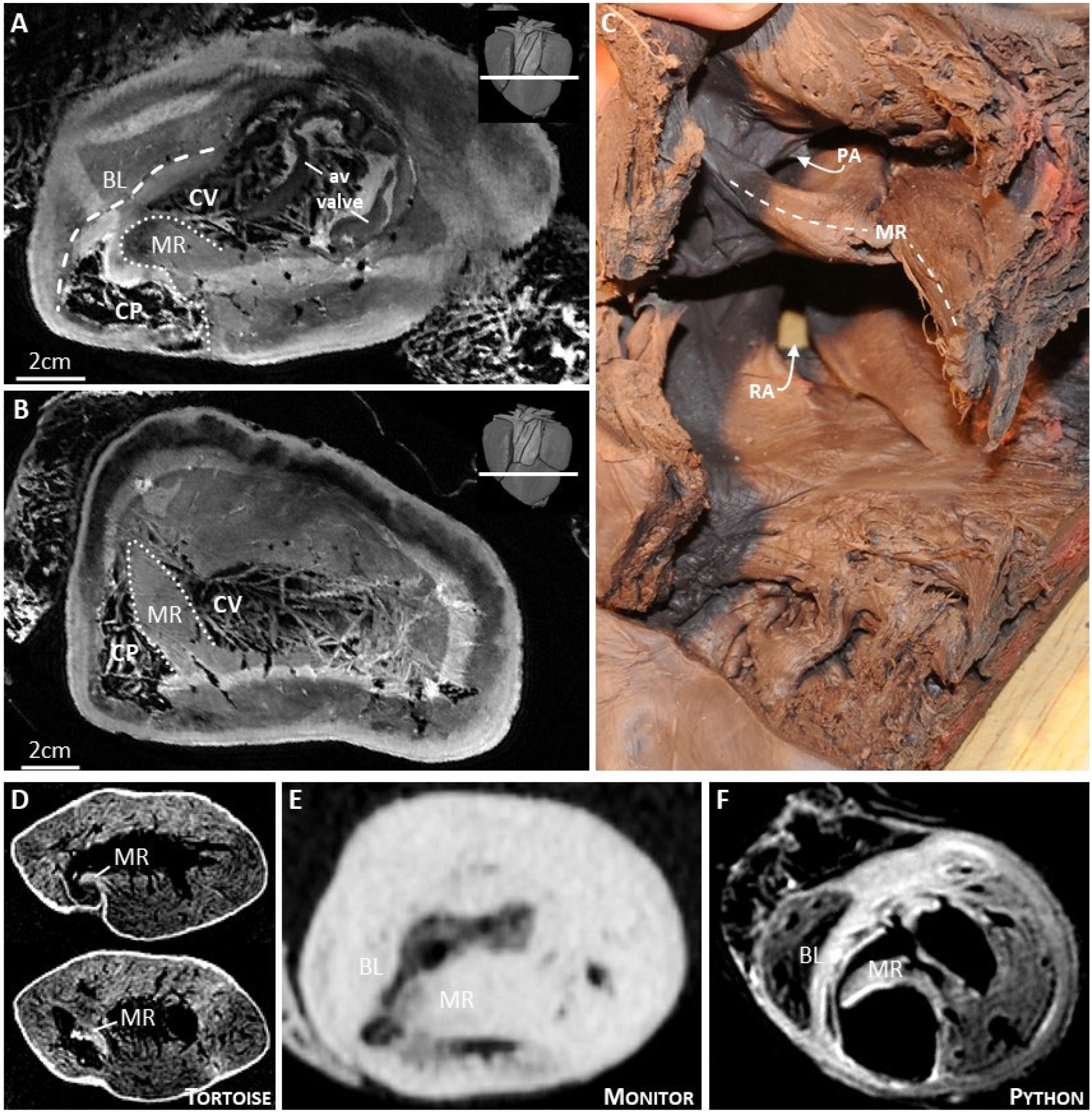
The muscular ridge of the ventricle. **A**. In the leatherback ventricle, the most pronounced septal structure is the muscular ridge (MR) opposite which is the bulbuslamelle (BL). These two structures mark the boundary between the cavum venosum (CV) and cavum pulmonale (CP). In the insert on the top-right, the section plane is indicated by a white line. **B**. The free edge of the muscular ridge has a spiral course and towards the apex it merges with myocardium of the dorsal-right wall. **C**. Muscular ridge of the Canadian heart, notice the large space between the free edge of the muscular ridge and the opposite wall. **D**. Muscular ridge of the tortoise. **E**. The muscular ridge of the monitor is exceptionally large. **F**. In the python, the muscular ridge and the bulbuslamelle are both exceptionally well-developed.

A single truncus arteriosus harbored the left and right aorta and the pulmonary artery in the specimens from Ballum and Canadian, as is typical in reptiles (Figs. 2C, 3C). In the Ballum heart, we surveyed the entire truncus in three planes for foramens between the major arteries. Only one potential foramen was found (Fig. 6). It was found in the shared wall of the left and right aorta, but even if it was patent it was tiny. The entire truncus was dissected in the Canadian heart without any foramen or large dimples being identified in or between any vessels.

**Figure 6.**
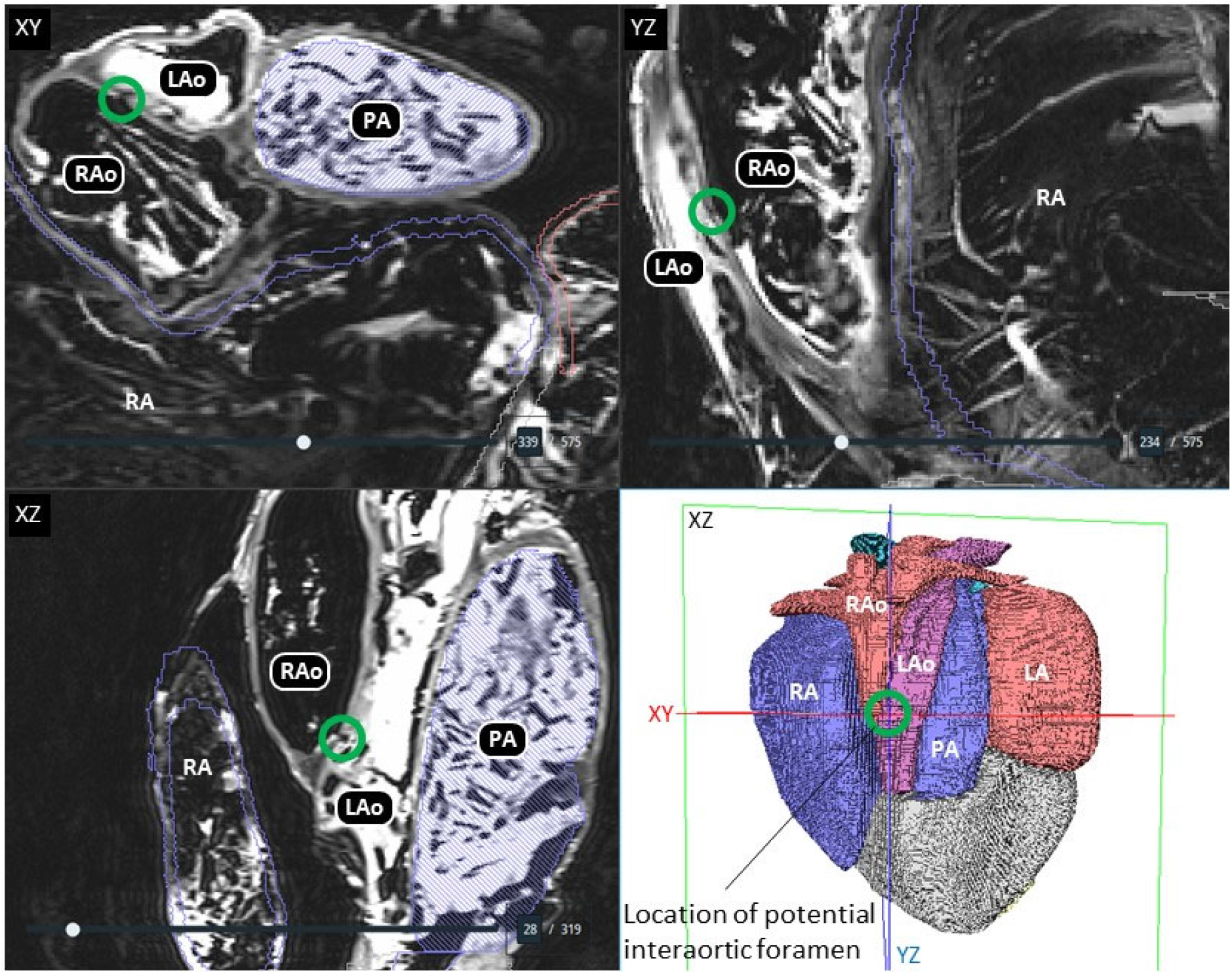
No pronounced interarterial foramens. In the Ballum turtle, we investigated all images in the three orthogonal planes (XY, YZ, XZ) but we did find such foramens as described by Adams (1962). A potential but very small foramen (green ring) was found at approximately the position where Adams (1962) describes one foramen.

## Discussion

Our assessment indicates that the leatherback turtle heart resemble that of other chelonians and does not exhibit obvious specializations for functional separation of the ventricle as it has been demonstrated in monitors and pythons. Overall, our description supports O’Donoghue’s original conclusion that the leatherback heart is “typically” chelonian with an internal anatomy “much like” that of the green sea turtle (O’Donoghue, 1918). Leatherback turtles are huge, with adults weighing hundreds of kilograms and even approaching a ton (Eckert & Luginbuhl, 1988; Georges & Fossette, 2006), and if the heart has exceptional capacities, they are achieved with a large but typical chelonian heart. We did not identify any large interarterial foramens such as those reported from the specimen examined by Adams (1962). We found no foramen in the Canadian specimen, but in the Ballum specimen, we did find a potential foramen in the shared wall of the left and right aortas, but it was so small that it would not have qualified for surgical repair had it been a human atrial septal defect (Webb & Gatzoulis, 2006).

Several anatomical traits of the leatherback heart resemble the typical chelonian heart and not those of monitors and pythons. The position of the coronary vessels on the ventricular surface are not consistent with ventricular septation as in monitors and pythons (or crocodylians) (Jensen *et al*., 2014). The leatherback ventricular apex is anchored by a gubernaculum cordis, which is a common trait in chelonians and lizards, but not in monitors and snakes (MacKinnon & Heatwole 1981; Jensen *et al*., 2014). There is no vertical septum that extends to the vicinity of the atrioventricular valve. Compared to monitors and pythons, the muscular ridge is small. It has previously been stated that amongst sea turtles, leatherbacks have a particularly well-developed muscular ridge, but detailed information was not provided (Wyneken 2001, 2009). We have previously quantified the size of the muscular ridge in reptile ventricles from cross sections similar to Figure 5A (Jensen *et al*., 2014) and we find the Ballum specimen to a have a typical rather than enlarged muscular ridge.

A particularly well-developed trait in monitors and pythons is a bi-layered configuration of the muscular ridge and the bulbuslamelle (Jensen *et al*., 2014), best observed in a dilated ventricle. Since the Ballum heart was contracted, we could not determine whether the muscular ridge and the bulbuslamelle had a similar bi-layered configuration. In monitors and pythons, there are also cushions of mesenchymal tissue, where the muscular ridge and the bulbuslamelle abut each other (Jensen *et al*., 2010b; Hanemaaijer *et al*., 2019). We did not observe similar cushions in the leatherback hearts, but the absence of evidence is obviously not solid evidence of absence.

Our description of a typical chelonian heart in the leatherbacks strongly indicate a considerable capacity for cardiac shunting and a lack of ventricular pressure separation. Other chelonians have large cardiac right-to-left shunts during diving (i.e. where oxygen-poor venous blood entering the ventricle from the right atrium is delivered to the aortas (e.g. Wang et al., 1997; Hicks, 1998). It is possible that leatherbacks also have large right-to-left cardiac shunt when diving, although the functional role of this shunt pattern remains to be established in diving reptiles (Hicks and Wang, 2012; Burggren et al., 2020). Cardiac right-to-left shunts lead to reductions in the arterial oxygen levels (i.e. both the partial pressure of oxygen and the oxygen concentration) and it has therefore been argued that functional division of the ventricle, as seen in monitors and pythons, provides for more efficient oxygen delivery and would be required for the ability to maintain the high aerobic metabolism needed for sustained physical activity. The leatherback turtle has some degree of endothermy and performs long migrations, so it would be expected that this species, in particular, could have evolved a more divided ventricle. Our present study shows that this is not the case. As an alternative explanation, it is possible that the leatherback fuel the high aerobic metabolism during migration by virtue of a large stroke volume provided by the very large heart. The relative large size of the heart has to be confirmed in additional specimens and the inferred large stroke volume would have to be validated by measurement of arterial blood flows in live animals.

Most chelonian hearts have a cartilage or bone deposit where the muscular ridge merges with the truncus arteriosus (O’Donoghue, 1918, Lopez *et al*., 2003; Poelmann *et al*., 2017). We saw a patch of different signal intensity on the MRI at this location, but we could not ascertain that this was indeed cartilage on the basis of the CT. We also saw straight lines in the ventricular cavities, which obscured the region under the atrioventricular valve where strands of connective tissue have been reported in several chelonian species (Greil, 1903; Acolat, 1943; Jensen *et al*., 2014). Such fine strands are not only easily missed, but they are perhaps lost during dissection when coagulated blood is removed from the ventricle. Adams (1962) describes numerous dimples on the luminal side of the arterial walls of his single specimen. We did not detect such dimples in either the Ballum or Canadian hearts.

In conclusion, despite leatherback turtles exhibiting adaptations to their unusual lifestyle, such as a high hematocrit (Lutcavage *et al*., 1990), the heart itself is typically chelonian in most regards. The only exceptional cardiovascular trait may be the relatively large heart that may provide for large stroke volumes.

## Acknowledgements

This work was supported by the Lundbeck Foundation (Grant# R324-2019-1470). A special thanks to Charlotte Bie Thøstesen for inviting TW and BJ to join the dissection of the Ballum leatherback and also thanks to the other members of the dissection team; Aage Kristian Olsen Alstrup, Daniel Klingberg Johansson, Hans Viborg Kristensen, Michael Paul Jensen, and Tim Kåre Jensen. Rasmus Buchanan and Jaco Hagoort gave technical assistance.

## Data Availability Statement

The data that support the findings of this study are available from the corresponding author upon reasonable request.

