## Supplementary Figure 1 for "Anatomy of the heart of the leatherback turtle"

### Information on the use of this interactive 3D-PDF

**All structures**
Sinus venosus
Sinuatrial valve
Right atrium
Atrial septum
Lung vein
Left atrium
Atrioventricular valve
**Ventricle**
- Ventricular wall
- Muscular ridge
- Bulbuslamelle
- Cavum arteriosum
- Cavum venosum
- Cavum pulmonale

Right aorta
Left aorta
Pulmonary artery
Gubernaculum cordis
Bronchi
Esophagus
Esophageal spines

Selected structure:

Amsterdam UMC
University Medical Centers

Department of Medical Biology

#### Selection of structures

The top left panel contains buttons to show or hide (groups of) structures, or to make them transparent.

**button example:** show / transparent / hide — Right atrium

After a single click on a 3D structure, the structure will be highlighted and the name of the structure will appear below "Selected structure". With the buttons next to the structure name, the appearance of this structure can be changed. Clicking next to the 3D object will deselect the structure.
For more advanced selection options, right-click on the 3D model and choose: “Show Model Tree”.

#### The intended use of this model

This model is intended to be used to better understand the approximate shape and position of major structures and cavities of the leatherback sea turtle heart.

#### Limitations of this model

Some structures are incompletely reconstructed because their full extent could not be recognized;
### Sinus venosus
### Lung vein
### Sinuatrial valve
There is an artificially sharp border between the ventricular muscle and;
### Muscular ridge
### Bulbuslamelle
There is an artificially sharp border between;
### Cavum arteriosum and Cavum venosum
### Cavum venosum and Cavum pulmonale
The trabecular structure of the atrial and ventricular walls is not reconstructed. One consequence is that the cavums appear to be very small cavities. In the actual ventricle, the wall is spongy and thus both wall and cavity and the cavums extend almost to the epicardium.

#### Technical Notes

View this PDF file in a recent version of Adobe Acrobat Reader: https://get.adobe.com/nl/reader/

3D interaction is only possible on MS Windows or Mac OS. Javascript and playing of 3D content must be enabled.

*Edit, Preferences* to ensure the following:

1) In *JavaScript*
   - enable *Enable Acrobat JavaScript*
2) In *Multimedia &3D*
   - enable *Enable playing of 3D content*
   - disable *Show 3D Orientation Axis*
   - *Optimization Scheme for Low Framerate: None*

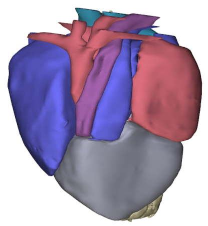
Ventral

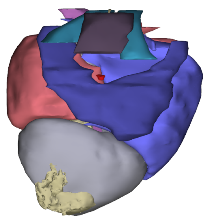
Dorsal

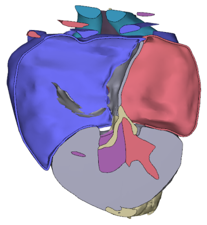
3 chamber view

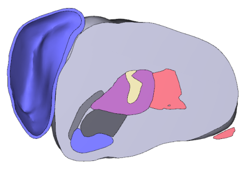
Transverse (muscular ridge)

### Heart of the leatherback sea turtle

**All structures**
- Sinus venosus
- Sinuatrial valve
- Right atrium
- Atrial septum
- Lung vein
- Left atrium
- Atrioventricular valve

**Ventricle**
- Ventricular wall
- Muscular ridge
- Bulbuslamelle
- Cavum arteriosum
- Cavum venosum
- Cavum pulmonale

- Right aorta
- Left aorta
- Pulmonary artery
- Gubernaculum cordis
- Bronchi
- Esophagus
- Esophageal spines

Selected structure:

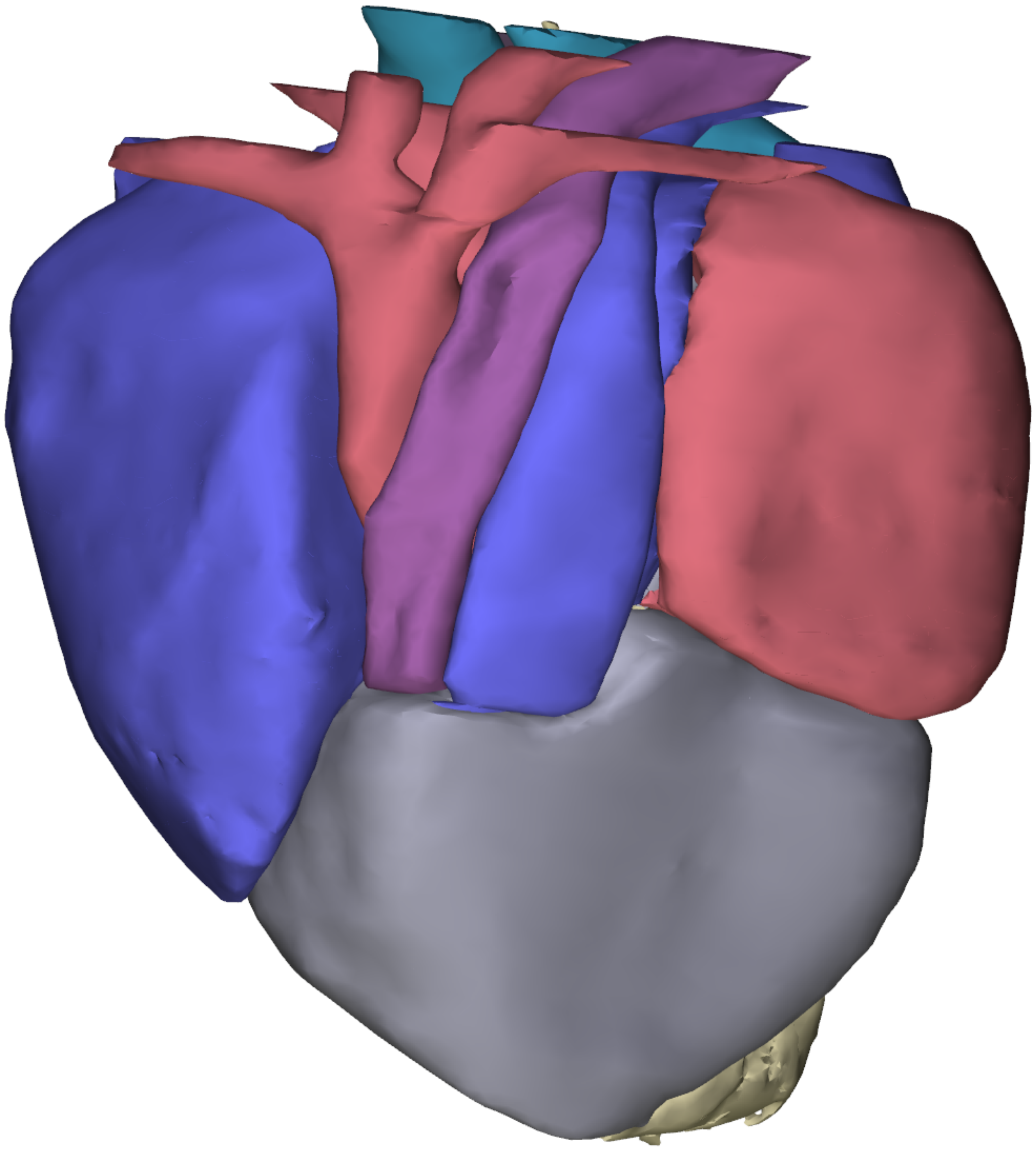

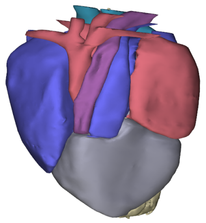

Ventral

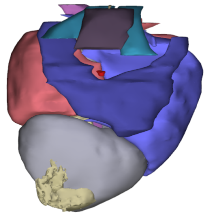

Dorsal

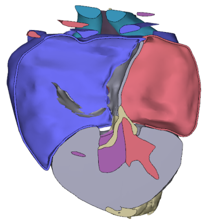

3 chamber view

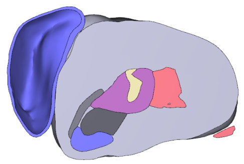

Transverse (muscular ridge)

Amsterdam UMC
University Medical Centers

Department of Medical Biology
